# Boid Inclusion Body Disease (BIBD) Is Also a Disease of Wild Boa Constrictors

**DOI:** 10.1101/2022.04.25.489483

**Authors:** Alejandro Alfaro-Alarcón, Udo Hetzel, Teemu Smura, Francesca Baggio, Juan Alberto Morales, Anja Kipar, Jussi Hepojoki

**Author notes:** Corresponding author: Present address: University of Helsinki, Medicum, Department of Virology, Haartmaninkatu 3, FI-00290 Helsinki Finland, or. The authors contributed equally to this manuscript (Alejandro Alfaro-Alarcón, Udo Hetzel).

## Abstract

Reptarenaviruses cause Boid Inclusion Body Disease (BIBD), a potentially fatal disease, occurring in captive constrictor snakes boas and pythons worldwide. Classical BIBD, characterized by the formation of pathognomonic cytoplasmic inclusion bodies (IBs), occurs mainly in boas, whereas in pythons, for example, reptarenavirus infection most often manifests as central nervous system signs with limited IB formation. The natural hosts of reptarenaviruses are unknown, although wild constrictor snakes are among the suspects. Here, we report BIBD with reptarenavirus infection in indigenous captive and wild boid snakes in Costa Rica using histology, immunohistology, transmission electron microscopy, next-generation sequencing (NGS), and RT-PCR. The snakes studied represented diagnostic post mortem cases of captive and wild caught snakes since 1989. The results from NGS on archival paraffin blocks confirm that reptarenaviruses have been present in wild boa constrictors in Costa Rica already in the 1980s. Continuous sequences de novo assembled from the low-quality RNA obtained from paraffin embedded tissue allowed the identification of a distinct pair of reptrarenavirus S and L segments in all studied animals; in most cases reference assembly could recover almost complete segments. Sampling of three prospective cases in 2018 allowed examination of fresh blood or tissues, and resulted in identification of additional reptarenavirus segments and hartmanivirus co-infection. Our results show that BIBD is not only a disease of captive snakes, but also occurs in indigenous wild constrictor snakes in Costa Rica, suggesting boa constrictors to play a role in natural reptarenavirus circulation.

**IMPORTANCE:** The literature describes cases of boid inclusion body disease (BIBD) in captive snakes since the 1970s, and in the 2010s, others and we identified reptarenaviruses as the causative agent. BIBD affects captive snakes globally, but the origin and the natural host of reptarenaviruses remains unknown. In this report, we show BIBD and reptarenavirus infections in two native Costa Rican constrictor snake species, and by studying archival samples, we show that both the viruses and the disease have been present in wild snakes in Costa Rica at least since the 1980s. The diagnosis of BIBD in wild boa constrictors suggests that this species plays a role in the circulation of reptarenaviruses. Additional sample collection and analysis would help to clarify this role further and the possibility of e.g. vector transmission from an arthropod host.

## INTRODUCTION

The first reports on boid inclusion body disease (BIBD) in captive snakes, mainly affecting members of the families *Boidae* and *Pythonidae*, go back to the 1970s (1). Others and we identified novel arenaviruses as the causative agent(s) in the early 2010s (2–8), and in 2015, the BIBD associated arenaviruses formed the genus *Reptarenavirus* in the family *Arenaviridae*, consequently leading to the formation of the genus *Mammarenavirus* (the previously known arenaviruses) (9). Up to now, studies have described BIBD only in captive snakes, and the origin of reptarenaviruses remains unknown. However, our group recently confirmed the disease also in native captive boa constrictors in Brazil, providing evidence that reptarenaviruses may circulate in indigenous snake populations (10). As the name implies, BIBD manifests by the formation of eosinophilic and electron-dense cytoplasmic inclusion bodies (IBs) within almost all cell types (1, 11, 12). In fact, the ante mortem diagnosis of BIBD relies on the detection of IBs in cells, for example blood cells in cytological specimens of blood smears, or hepatocytes in liver biopsies (1, 13). More recently, the identification of reptarenavirus nucleoprotein (NP) as the main component of the IBs has allowed the use of immunohistochemistry to support the diagnosis, significantly increasing the diagnostic specificity and sensitivity (14, 15). Due to reasons unknown, IBs appear to be more common in reptarenavirus-infected boas than pythons (2, 4, 7, 8, 13, 15). In naturally infected pythons, reptarenavirus infection predominantly manifested as central nervous system (CNS) signs which include regurgitation, “star-gazing”, head tremors, disorientation, and “corkscrewing”; these were also described in early reports of boas with BIBD but seem not to occur more recently (1). Indeed, experimental infection of pythons with reptarenavirus isolates induced CNS signs whereas the infected boas remained without overt clinical signs (7, 8). In fact, different from the early reports on BIBD, the majority of boa constrictors with BIBD nowadays appear clinically healthy (15–17).

Boas and pythons are non-venomous constrictor snakes inhabiting biotopes of the neotropics and tropics. Boas occur in Central and South America and Madagascar, i.e. in the New World, whereas pythons occupy habitats in Africa, Asia and Australia, i.e. in the Old World. Thus, the natural habitats of boas and pythons do not overlap much (The Reptile Database, http://reptile-database.org/). The literature has reported several of the more than 100 known boa and python species to be susceptible to BIBD (1).

Costa Rica harbours four indigenous boid species, *Boa constrictor* (LINNAEUS, 1758), *Corallus annulatus* (COPE, 1875), *Corallus ruschenbergerii* (COPE, 1875) and *Epicrates cenchria* (LINNAEUS, 1758), distributed in different, partly overlapping habitats. Of these, *B. constrictor*, *C. annulatus* and *E. cenchria* have been shown to be susceptible to BIBD (1, 15, 18). *B. constrictor* has a wide distribution ranging from Florida, US (introduced species), to South America. In Costa Rica, it is distributed all over the country, inhabiting tropical and subtropical rain forests, versants and Valle Central, at 0 – 1500 metres above sea level. The nocturnal species lives mostly close to the ground and is often found close to coffee plantations, predominantly preying on mammals and birds (19). *C. annulatus* and *C. ruschenbergii* are arboricole snakes. The former is widely distributed in Central America and Colombia. In Costa Rica it inhabits the canopy of the Carribean subtropical and tropical rain forest, 0 – 650 metres above sea level (19, 20). *C. ruschenbergii* occupies habitats of the tropical rain forest of the Pacific south, i.e. Costa Rica, Panama, northern Colombia, northern Venezuela, Trinidad and Tobago, and Isla Margarita, at 0 - 500 metres above sea level (19, 20). The nocturnal *E. cenchria* has a wide distribution in Central and South America, from Costa Rica to Argentina, feeding predominantly on bats and rodents (21). In Costa Rica it inhabits tropical rain forests, the Carribean versant, Pacific northwest and Pacific south, at altitudes between 0 - 500 metres above sea level (19).

The family Arenaviridae currently comprises four genera, *Mammarenavirus* (rodent- and bat borne viruses), *Reptarenaviruses* (the BIBD associated viruses), *Hartmaniviruses* (found in boa constrictor), and *Antennavirus* (fish viruses) (22, 23). The genome of reptarenaviruses is a bisegmented negative-sense RNA with ambisense coding strategy (22). The S segment encodes glycoprotein precursor (GPC) and nucleoprotein (NP), and the L segment gives rise to zinc finger matrix protein (ZP) and RNA-dependent RNA polymerase (RdRp) (22). We identified Haartman Institute Snake virus-1 (HISV-1) during a sequencing study of snakes with BIBD (6), which after further characterization (24) became the founding member of the genus *Hartmanivirus* (25). HISV-1 and all other known hartmaniviruses, lack the ZP gene (24) and do not appear to contribute to BIBD pathogenesis although they often co-exist with reptarenaviruses in snakes with BIBD (16, 24).

The literature best describes the occurrence of BIBD in captive snake collections in Europe, North America, Asia and Australia (14-17, 26, 27). We recently reported BIBD and reptarenavirus infection in captive boa constrictors native to Brazil, however, the lack of detailed information on the origins of the studied snakes and their potential contacts during captivity did not allow us to conclude that reptarenaviruses are present in wild snakes (10). Here, by studying captive and wild-caught native wild snakes with BIBD from Costa Rica, we show that BIBD occurs in wild boa constrictors, in conjunction with reptarenaviruses and hartmaniviruses.

## MATERIALS AND METHODS

### Animals

The studied animals included 11 snakes submitted to the Department of Veterinary Pathology, Universidad Nacional, Heredia, Costa Rica, for diagnostic post mortem examination. Among these were six captive snakes, four *B. constrictor* and two *C. annulatus*, examined in 2004 and 2006 (animals C1-C6), and five wild boa constrictors that were caught and immediately euthanised due to severe clinical disease in 1989, 2012 and 2018 (animals W1–W5) (Table 1). Due to the unclear taxonomic classification of *Boa constrictor* subspecies (28, 29), we only refer to *Boa constrictor*. From all animals, specimens from selected organs and tissues were collected at necropsy and were fixed in 10% buffered formalin for histological examination. From the snakes examined in 2018, additional tissue samples (brain, liver) were collected and frozen at −80 °C for virus isolation and RNA extraction.

**Table 1:**
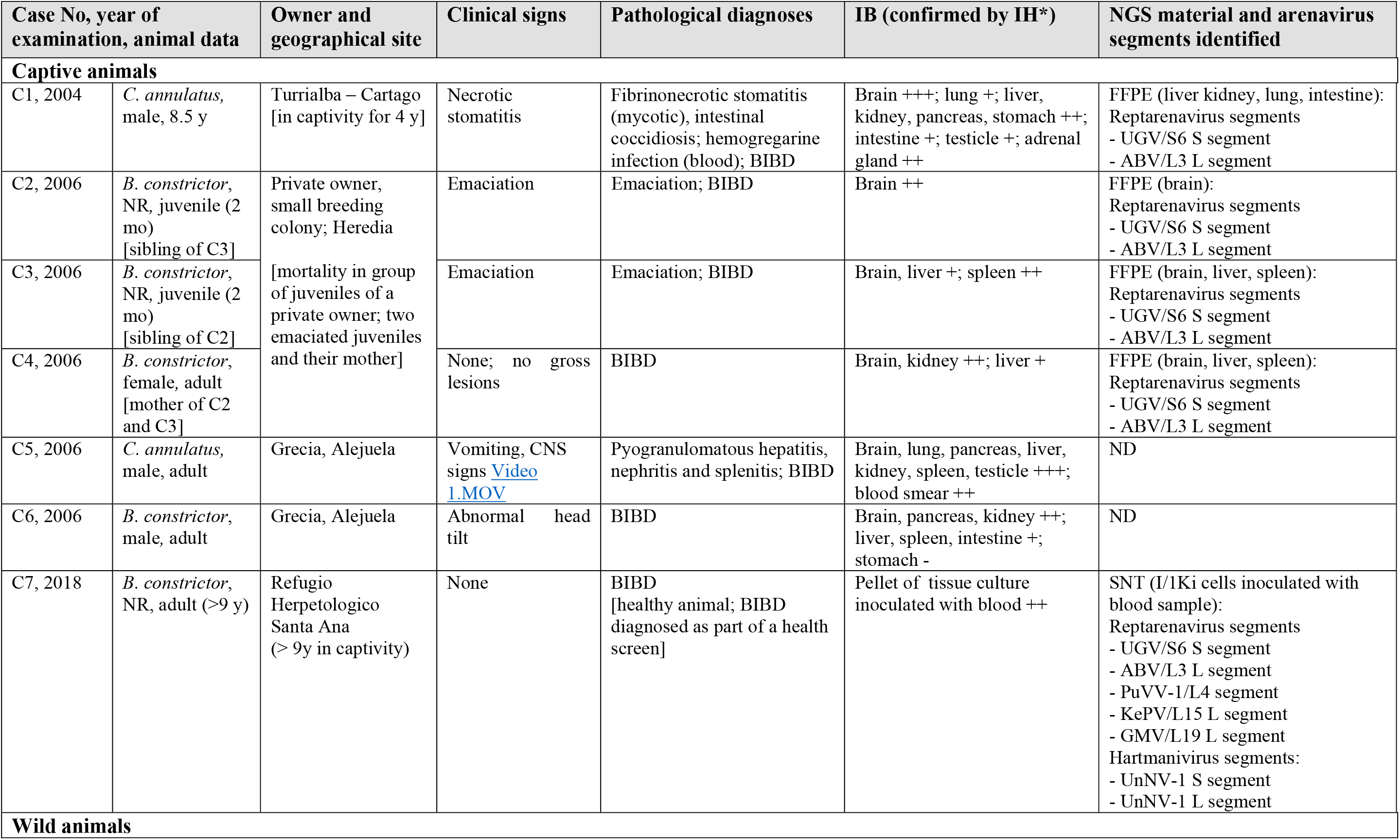
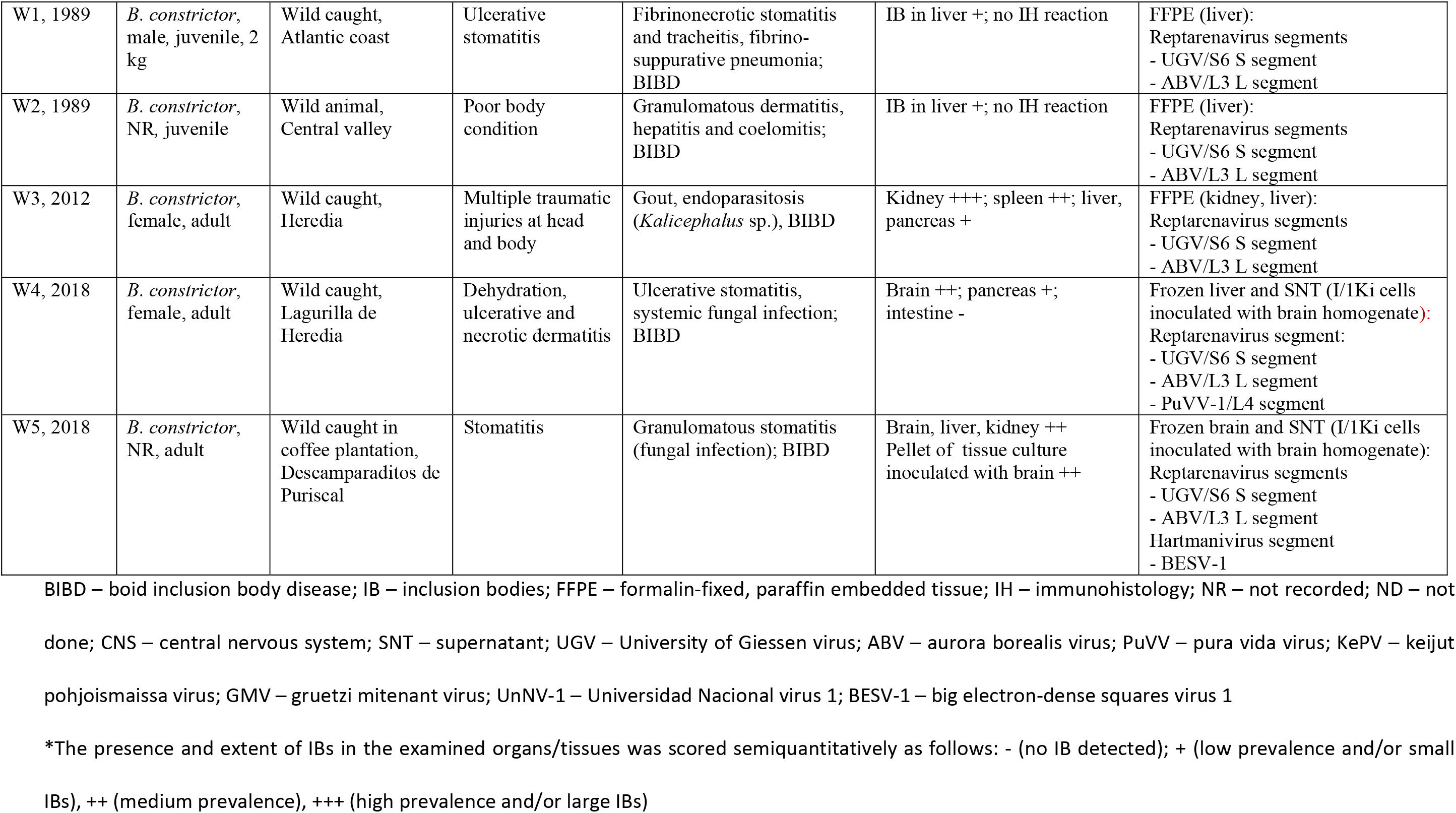
Animals included into the study, their origin, clinical signs and pathological diagnoses, detection of reptarenavirus and hartmanivirus nucleoprotein in tissues or a blood inoculated tissue culture, and reptarena- and hartmanivirus identification through NGS.

The study also included one apparently healthy *B. constrictor* from a reptile sanctuary that the owners wished to be screened for BIBD and hemoparasites before moving it to housing with closer contact to other snakes. From this animal (1.7), a blood sample was collected in a 1.3 ml K3E EDTA tube (Sarstedt) by venipuncture of the caudal tail vein, aliquoted (one sample diluted 1:1 in RNAlater (ThermoFisher Scientific), one direct) and frozen at −80 °C for RNA extraction and virus isolation, respectively. In addition, blood smears were prepared.

No ethical permissions were required for these diagnosis-motivated blood and tissue samplings, which were requested by the owners and permitted by the authorities, respectively. To specify, all post mortem examinations of wild snakes were conducted with approval by the Ministry of Environment and Energy (MINAE) (wildlife authority) through permit R-SINAC-PNI-ACLAC-039, and with the support of the animal health authority, the National Animal Health Service through the permit SENASA-DG-0277-18.

### Histology and immunohistology, cytology

Formalin-fixed tissue specimens were trimmed and routinely paraffin wax embedded. Sections (3-5 µm) were prepared at the time of diagnosis and stained with hematoxylin-eosin (HE). At a later stage, in 2021, consecutive sections were subjected to immunohistological staining for reptarenavirus NP as described (4, 10). For cytological examination of blood cells for the presence of inclusion bodies, the current standard ante-mortem diagnostic approach (1, 15), blood smears taken from animals C1, C5 and C7 were stained with May-Grünwald-Giemsa and examined by light microscopy as described (16, 30).

### Cell culture and virus isolation

The *Boa constrictor* kidney derived cell line, I/1Ki, maintained at 30 °C, 5% CO_2_, was used for virus isolation attempts as described (4). Tissue cubicles of 1-2 mm^3^ of liver (animal 2.4) and brain (animals 2.5) were defrosted, mechanically homogenized in 6-8 ml of fully supplemented growth medium (Minimal Essential Medium Eagle [Sigma], 10% of fetal bovine serum [Gibco], 10% of tryptose-phosphate broth [Sigma-Aldrich], 2 mM L-glutamine 1.4 mM Hepes buffer, 100 IU/ml gentamycin), spun for 5 min at 300 RCF and filtered through a 0.45 µm syringe filter (Millipore). Confluent I/1 Ki cells (75 cm^2^ culture flasks) were inoculated with 1 ml of filtered inoculum for 6-12 h, followed by a media exchange.

For virus isolation from the blood (animal C7), we inoculated a sub-confluent culture of I/1Ki cells with 300 µl of blood diluted in 3 ml fully supplemented growth medium filtered through a 0.45 µm syringe filter. After 4 h adsorption at 30°C, 5% CO_2_, 4 ml of fresh medium was added and the culture incubated overnight at 30°C, 5% CO_2_. The following day, we washed the cells once with fresh medium and continued the incubation at 30°C, 5% CO_2_.

For all cultures, the cells were kept for 6 days and the media exchanged after 3 days. Supernatants from both time points were collected, mixed and frozen at −80 °C. For the preparation of cell pellets from the inoculated cell cultures, cells were trypsinised (Trypsin 10x, Merck Biochrome) for 5 min at room temperature, then spun at 1,000 x RCF for 3 min. The resulting pellet was either fixed in 4% buffered paraformaldehyde overnight and routinely embedded in paraffin wax for histology and immunohistology, or was fixed in 2.5% phosphate buffered glutaraldehyde and routinely epoxy resin embedded for transmission electron microscopy, as previously described (24, 31).

### Next generation sequencing (NGS) and genome assembly

RNAs were extracted from paraffin embedded tissue samples of animals C1-C4 and W1-W3 (Table 1) using the RNeasy FFPE Kit (QIAGEN) according to manufacturer’s protocol. The RNA extraction from other sample materials, including the blood of animal C7 (direct from 300 μl blood frozen with an equal amount of RNAlater), followed the Trizol reagent (Roche) protocol as described (10, 30). NGS library preparation from isolated RNAs, sequencing, and subsequent genome assembly was done as described (10, 30, 32).

### Phylogenetic analyses

The amino acid sequences of previously identified hartmaniviruses were retrieved from GenBank. MAFFT E-INS-i algorithm (33) was used for aligning the sequences with those of the viruses identified in this study. Bayesian Monte Carlo Markov chain (MCMC) method implemented in MrBayes v3.2.6. (34) served for inferring the best-fit amino acid substitution models and constructing phylogenetic trees. In total 500,000 generations with sampling at every 5,000 steps were run in MrBayes. The final standard deviations between two runs were <0.02 for all analyses.

## RESULTS

### Case descriptions

In 2004 – 2006, we examined six captive snakes, two *Corallus annulatus* and four boa constrictors (Table 1, animals C1-C6,). The first (animal C1), a male *C. annulatus*, had been wild caught and kept in captivity for 4 years, housed with another snake of the same species, before falling ill in 2004. It was euthanized due to necrotic lesions in the oral cavity. The post mortem examination revealed a fibrinonecrotic stomatitis with intralesional fungi and bacteria (identified in a direct smear) that remained unidentified as a microbiological examination was not performed. The parasitological examination of gut content found *Eimeria sp*. oocysts, and the examination of a blood smear detected hemogregarine infected red blood cells. The histological examination confirmed the gross findings and revealed the typical inclusion bodies (IBs) in parenchymal cells in various organs, including neurons in the brain, tubular epithelial cells in the kidney, hepatocytes, airway and lung epithelial cells, pancreatic epithelial cells as well as gastric, small and large intestinal epithelia (Table 1, animal C1). The blood smear that had been prepared immediately after the animal’s death showed presence of IBs within erythrocytes.

In 2006, diagnostic post mortem examinations identified five cases of BIBD in captive snakes. The owner of a small private breeding colony had observed high mortality in a group of juvenile captive-bred boa constrictors (10 of 27 died), due to which he submitted two emaciated snakes from the group (animals C2 and C3) for euthanasia and diagnostic post mortem examination. The examination did not reveal any other gross changes. The histological changes were restricted to the presence of abundant cytoplasmic IBs in the examined organs, i.e. neurons and ependymal cells in the brain, hepatocytes, lymphocytes and erythrocytes in the spleen. After the BIBD diagnosis in the two juvenile animals and having been informed that reptarenavirus infection in the juveniles could be a consequence of vertical transmission from the parental animals (30), the owner requested euthanasia and a diagnostic post mortem examination of their mother (animal C4). Also in this animal, the pathological changes were restricted to the presence of cytoplasmic IBs in parenchymal cells of multiple tissues, brain, liver and kidney, thus confirming that the animal had also suffered from BIBD.

Another two adult snakes, a male *C. annulatus* (animal C5) and a male *B. constrictor* (animal C6) from a private breeder, were euthanized due to central nervous system signs (for video, see Table 1). The diagnostic post mortem examination of the *C. annulatus* revealed a pyogranulomatous hepatitis, nephritis and splenitis and cytoplasmic IBs in parenchymal cells of various organs, i.e. the brain, lung, pancreas, liver, kidney, spleen, and testicle and in blood cells (identified in a blood smear) (Fig. 1 and Table 1, animal C5) which also showed hemogregarine infection of red blood cells (Fig. 1E). The *B. constrictor* did not exhibit any gross or histological changes apart from the presence of abundant cytoplasmic IBs in parenchymal cells of brain, pancreas, kidney, liver, spleen and intestine (Table 1, animal C6).

**Figure 1.**
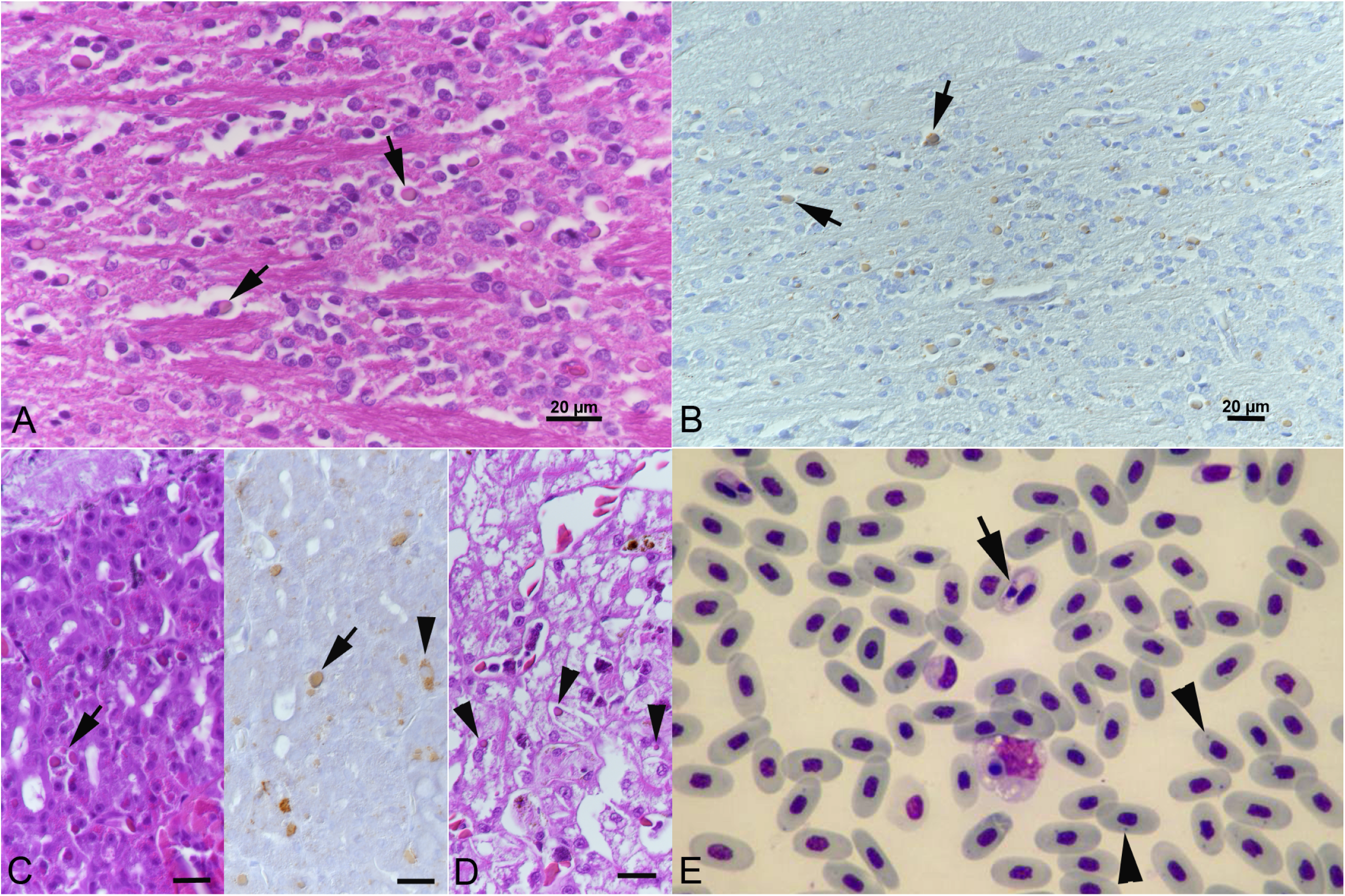
Animal C5; captive *C. hortulanus* with BIBD. **A, B**. Brain stem. Cytoplasmic inclusion bodies (IBs) (arrows) in neurons that are comprised of reptarenavirus nucleoprotein (B). **C**. Pancreas with IBs in epithelial cells (arrows). There are also individual cells with diffuse cytoplasmic viral antigen expression (right image, arrowhead). **D**. Liver with IBs in hepatocytes (arrowheads). **E**. Blood smear. Red blood cells with IBs (arrowheads) and hemogregarine infection (arrow). May-Grünwald-Giemsa stain. A, C (left), D: HE stain; B, C (right): immunohistology, haematoxylin counterstain. Bars = 20 µm.

The last animal of this series was a wild-caught *B. constrictor* (animal C7) that had been housed in captivity in a reptile park for over 9 years. The animal was one of several healthy snakes that underwent a health screen including an examination of a blood smear to determine the BIBD status and detect or exclude protozoan infection, as a basis for new co-housing plans. The animal exhibited the typical cytoplasmic IBs in multiple blood cells (mainly erythrocytes) leading to the diagnosis of BIBD but showed no evidence of protozoan infection. It was subsequently housed in an individual terrarium remote from other snakes and was then lost for follow-up investigations.

The remaining five snakes of the study represent wild boa constrictors that had been caught by the local authorities. All were then euthanized by a veterinarian due to severe clinical disease. The first two animals were examined in 1989. A juvenile *B. constrictor* was caught at the Atlantic coast (Table 1, animal W1) and found to suffer from a severe, focal extensive fibrinonecrotic and ulcerative stomatitis, a fibrinosuppurative tracheitis and a moderate multifocal fibrinosuppurative bronchopneumonia with multiple microabscesses. The bacteriological examination isolated *Escherichia coli* (not further specified) from the lesions, and the parasitological examination revealed a moderate intestinal endoparasitosis (nematodes, not further specified). The second animal (Table 1, animal W2) had been found with poor body condition in the central valley. It was transferred to the national zoological garden where it did not feed during the quarantine period, hence it was euthanized. The post mortem examination revealed a multifocal ulcerative and pyogranulomatous to necrotic dermatitis with intralesional fungal structures (fungal culture: *Fusarium spp*.) as well as a multifocal granulomatous coelomitis (cranially to kidneys) and hepatitis. The animal also suffered from a moderate intestinal endoparasitosis (nematodes, not further specified). In both animals, the histological examination of the liver revealed the typical cytoplasmic IBs within hepatocytes, leading to the diagnosis of BIBD.

The third animal (Table 1, animal W3) was caught in 2012. It was euthanized due to multiple traumatic injuries at head and body (Fig. 2A). The post mortem examination showed that the animal suffered from gout (Fig. 2B). The histological examination also revealed abundant cytoplasmic IBs in parenchymal cells of kidneys, spleen and liver.

In 2018, at a time when the group of investigators had started to collaborate to work up cases of BIBD in Costa Rica, another two wild caught sick boas (Table 1, animals W4 and W5) were submitted for a diagnostic post mortem examination. The first animal was euthanized due to a multifocal necrotic dermatitis (Fig. 2C) and was found in the post mortem examination to have suffered also from an ulcerative stomatitis as well as a multifocal to coalescing heterophilic and granulomatous hepatitis, nephritis and myocarditis with intralesional fungal structures (not further identified). The second animal was euthanized due to a granulomatous stomatitis with intralesional fungal structures (not further identified) without any other significant lesions. Both animals exhibited cytoplasmic IB in parenchymal cells in brain and pancreas and brain, liver and kidney, respectively (Fig. 3). In the brain of animal W5 these were also obvious as large, brightly eosinophilic partly square inclusions (Fig. 3A).

**Figure 2.**
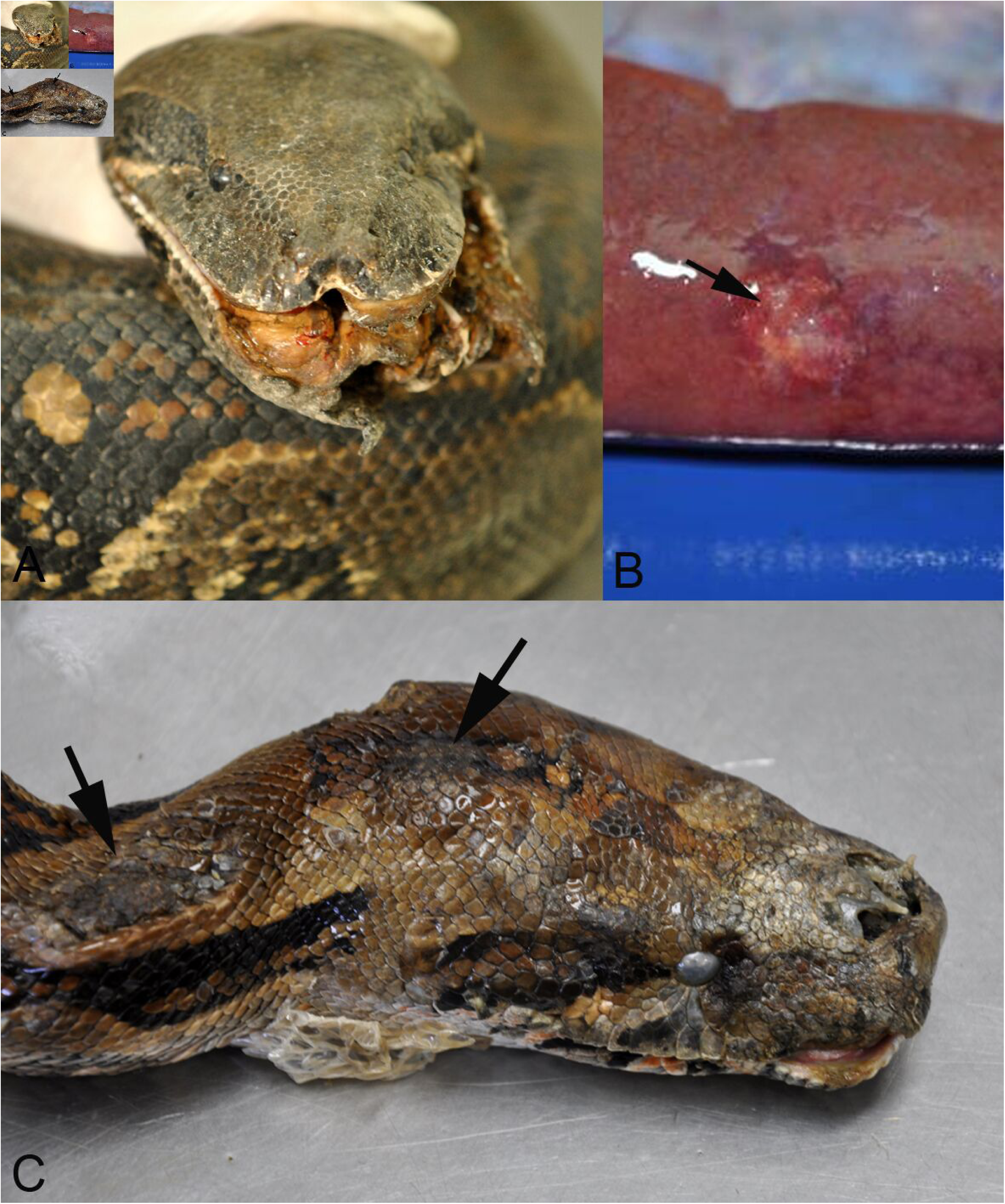
Gross findings in two wild caught boa constrictors that were euthanized due to incurable disease. **A, B**. Animal W3. **A**. Head with severe traumatic injuries. **B**. Liver with focal granulomatous inflammation in association with urate crystals (gout). **C**. Animal W5. Head with multifocal necrotic dermatitis (arrows).

**Figure 3.**
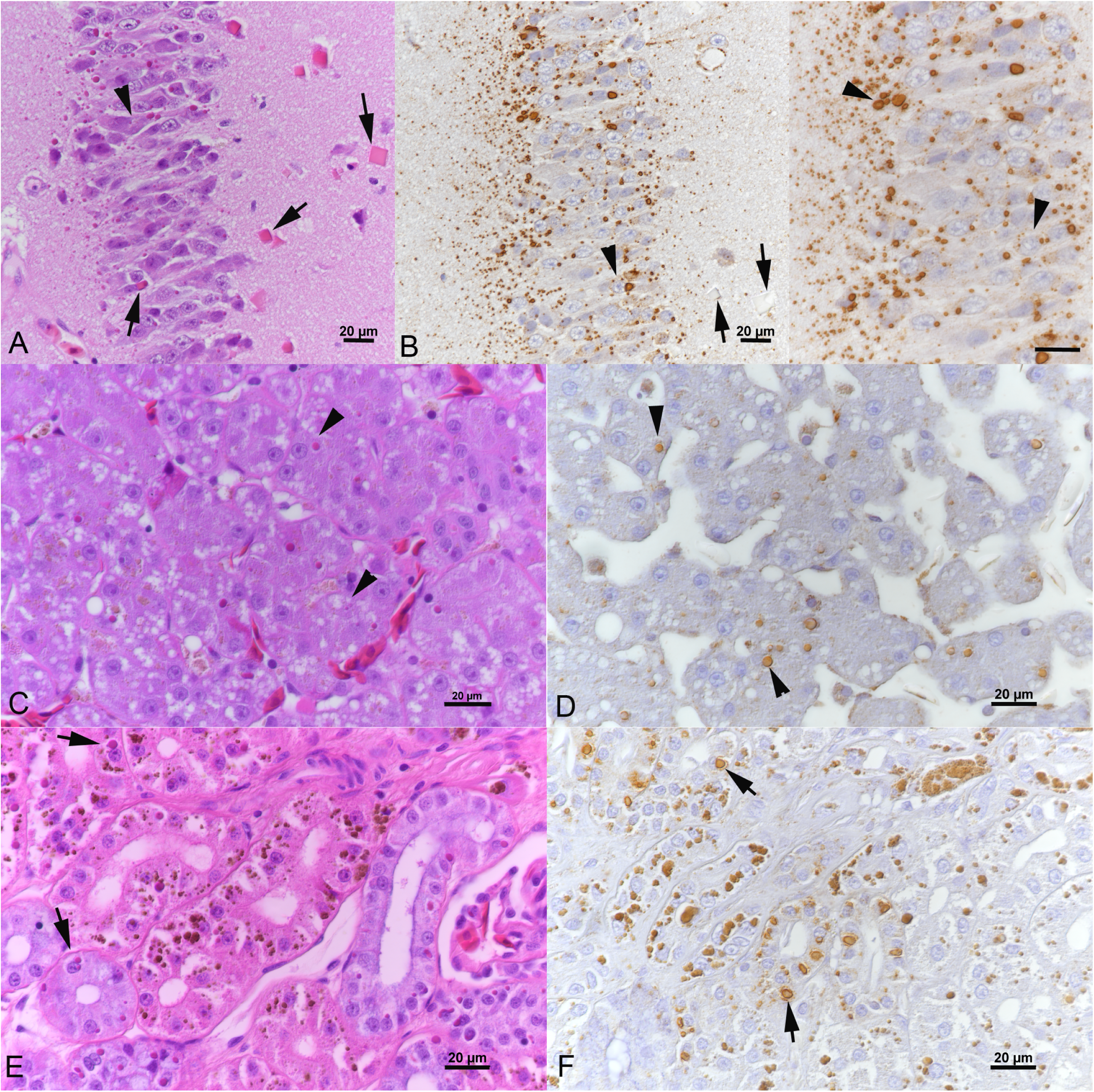
Wild caught boa constrictor (animal W5) with BIBD. Inclusion body (IB) formation and reptarenavirus nucleoprotein (NP) expression in tissues. **A, B**. Cerebral cortex with abundant cytoplasmic IBs in neurons. Some neurons exhibit more than one IB (arrowheads), others show large square IBs (A, arrows) in which reptarenavirus NP expression is restricted to a focal punctate reaction (B, left image, arrows). **C, D**. Liver with variably sized IBs in hepatocytes (arrowheads). **E, F**. Kidney with variably sized IBs in tubular epithelial cells (arrows). NB: The coarse brown pigment in the epithelial cells represents lipofuszin-like degradation products frequently observed in the kidney of boas. A, C, E: HE stain; B, D, F: immunohistology, haematoxylin counterstain. Bars = 20 µm.

### Confirmation of BIBD and reptarenavirus infection by immunohistology

In 2021, immunohistology for the detection of reptarenavirus NP was performed on tissue sections from all snakes. This yielded a positive result (Figs. 1B, C) thereby confirming reptarenavirus infection in all cases apart from the two wild snakes from 1989, the paraffin blocks of which had likely suffered from protein degradation during storage (Table 1).

### Virus isolation

Virus isolation was attempted from three boa constrictors examined in 2018 (Table 1). The blood from captive snake C7, a liver homogenate of wild caught snake W4, and a brain homogenate of wild caught snake W5 served to inoculate I/1Ki (*B. constrictor* kidney cell line) cell cultures. Immunohistology of cell pellets prepared after 6 days of inoculation showed extensive inclusion body formation and reptarenavirus NP expression in all inoculated cells (Fig. 4B). Hartmanivirus NP expression was not observed. Interestingly, like neurons in the brain of the affected snake, the cultures inoculated with the brain homogenate of animal W5 exhibited occasional large, brightly eosinophilic, square cytoplasmic inclusions (Fig. 4A) interpreted as overwhelming NP expression. Ultrastructurally, the cells were found to harbor the previously described early and late inclusions as well as dense, large square inclusions with a grid-like electron dense pattern (Fig. 4C-F) reminiscent of protein crystals.

**Figure 4.**
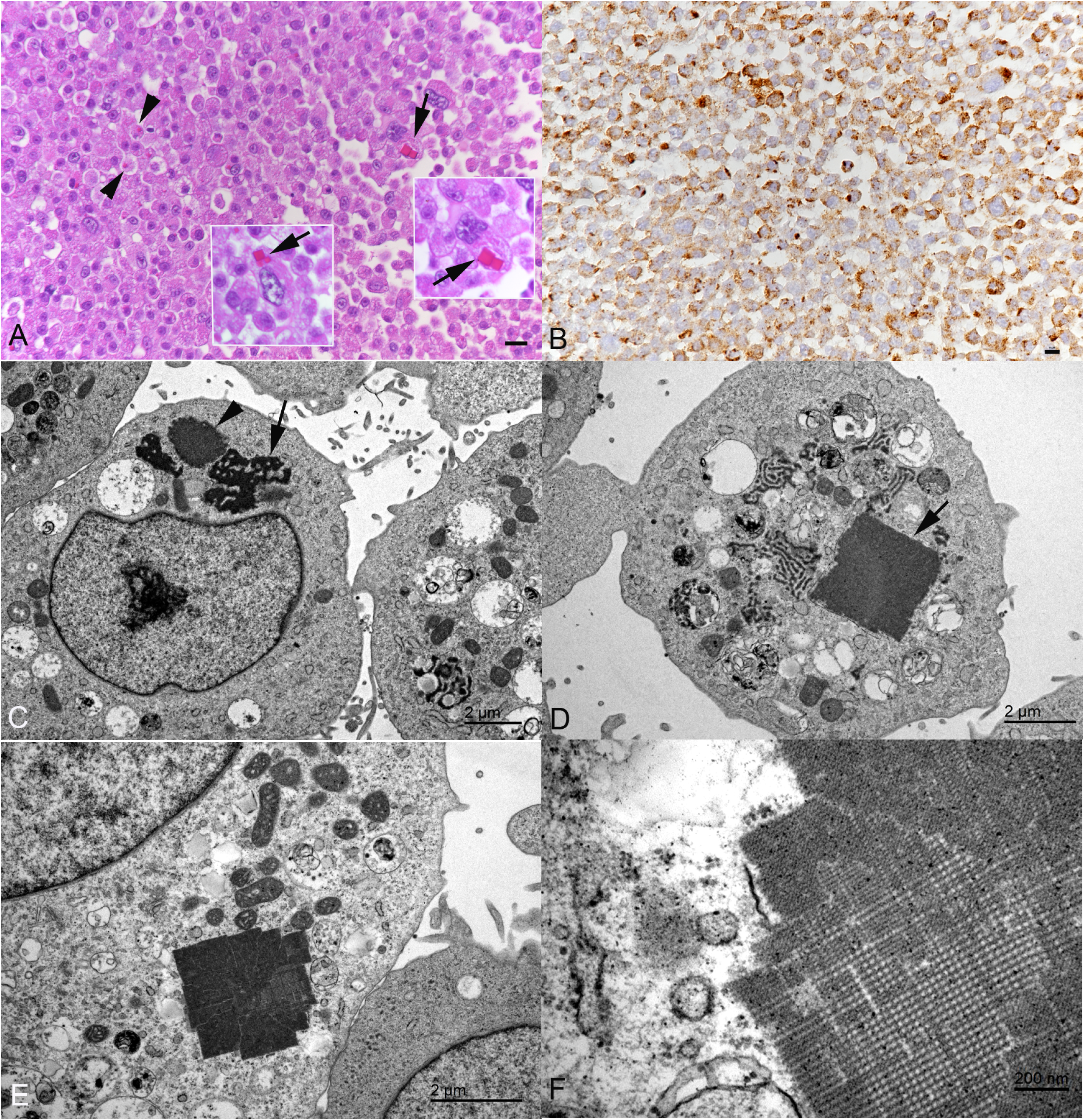
Wild caught boa constrictor (animal W5) with BIBD. Light and transmission electron microscopy findings in a cell pellet prepared from the *Boa constrictor* kidney derived cell line, I/1Ki, at 6 days post inoculation with brain homogenate. **A, B**. Inclusion body (IB) formation (A, arrowheads) and reptarenavirus nucleoprotein (NP) expression (B) in the cells. Like the neurons in the cerebral cortex (Fig. 3A), individual cells exhibit large square IBs (arrows). **C-F**. Ultrastructural features. The cells exhibit smaller and irregular early (arrow) and more complex older (arrowhead) IBs (C) and occasional large square IBs (D) with a grid like, directed structure (E, F).

### Identification of viruses

To confirm reptarenavirus infection and to identify the infecting viruses, we performed a metatranscriptomic analysis of the formalin-fixed paraffin embedded (FFPE) archival material (Table 1, animals C1-C4 and W1-W3), freshly frozen liver (Table 1, animal W4) and brain (Table 1, animal W5), and supernatants of cell cultures inoculated with tissue homogenates and blood (Table 1, animals C7, W4 and W5), respectively. In addition to the de novo genome assembly based approach utilized in our earlier studies (6, 10, 16, 24, 30, 32, 35), we used the lazypipe NGS pipeline for pathogen discovery (36). The lazypipe served to produce an overview of the microbial reads in the samples after removal of host genome from the dataset, demonstrating the presence of mainly viruses and bacteria (Fig. 5). Most of the eukaryotic reads remaining in the samples represented members of the families *Aspergillaceae* and *Thermoascaceae*, possibly representing the causative agent of the mycotic disease and/or reflecting contamination and deterioration of the archival paraffin blocks. Additionally, lazypipe identified some reads matching potential prey animals (families *Passerellidae*, *Sylviidae*, *Phasianidae* and *Columbidae*) and blood-sucking parasites (family *Argasidae* and *Trombidiidae*).

**Figure 5.**
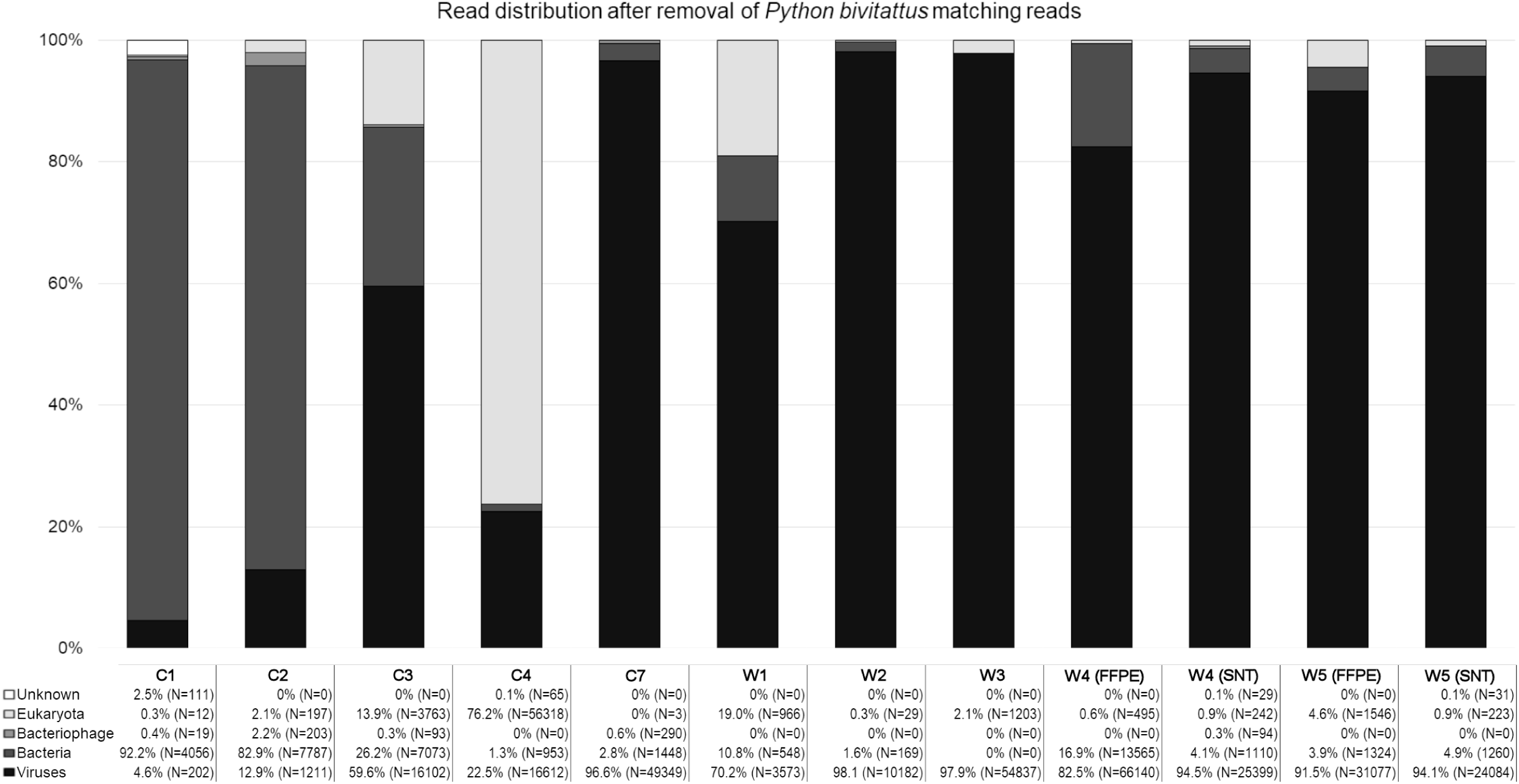
The distribution of reads in Lazypipe (36) analysis after the removal of reads matching to *Python bivitattus* genome. The y-axis represents percentage of reads. The reads matching viruses, bacteria, bacteriophages, eukaryota, and unknown (i.e. those not without match) are shown in colors ranging from black to white as indicated in bottom right corner. The number of reads matching each category are presented in written format (% of total reads of the sample and the number of reads in brackets) under each respective sample name.

The viruses identified through the lazypipe analysis of the samples that went through NGS are summarized in Table 2. The majority of the viral reads not matching arenaviruses included viruses (families *Partitiviridae*, *Chrysoviridae*, and *Botourmiaviridae*) infecting fungi, plants and possibly insects, but also viruses (families *Adenoviridae*, *Picobirnaviridae* and *Nodaviridae*) that may indeed have infected the studied animals. The analysis confirmed reptarenavirus infection in all of the studied animals, and identified hartmanivirus infection in two animals (Table 1, animals C7 and W5). For most samples, the standard lazypipe run did not produce complete segment-length contigs and we employed MIRA assembler (http://mira-assembler.sourceforge.net/docs/DefinitiveGuideToMIRA.html) as well as Bowtie2 (37) to assemble full length arenavirus S and L segments. As presented in Table 1, all animals studied carried the University of Giessen virus (UGV) S segment and aurora borealis virus 3 (ABV-3) L segment, i.e. S6 and L3 segments according to the nomenclature coined by Stenglein et al. (5). Two of the studied animals (Table 1, C7 and W4) showed presence of more than a single pair of reptarenavirus S and L segments, both harbored an L4 L segment earlier identified by Stenglein and colleagues (5), which we designated as L segment of pura vida virus 1 (PuVV-1). The analysis revealed two additional L segments, i.e. those of keijut pohjoismaissa virus 1 (KePV-1) and gruetzi mitenant virus 1 (GMV-1), respectively known as L15 and L19 following the Stenglein et al. nomenclature, in the animal C7. All of the identified reptarenavirus segments were highly similar to those described in earlier studies.

**Table 2.** Distribution of NGS reads matching to viruses as identified by Lazypipe (1).

| Animal no. (material) | No. Reads | No. Contigs | Virus family | Genus |
| --- | --- | --- | --- | --- |
| C1 (FFPE: liver, kidney, lung, intestine) | 195 | 3 | *Arenaviridae* | *Reptarenavirus* |
|  | 7 | 0 | *Picobirnaviridae* | unknown |
| C2 (FFPE: brain) | 1090 | 0 | *Partitiviridae* | unknown |
|  | 85 | 3 | *Adenoviridae* | *Mastadenovirus* |
|  | 32 | 1 | *Nodaviridae* | *Alphanodavirus* |
|  | 4 | 1 | *Arenaviridae* | *Reptarenavirus* |
| C3 (FFPE: brain, liver, spleen) | 15921 | 6 | *Arenaviridae* | *Reptarenavirus* |
|  | 64 | 0 | *Partitiviridae* | unknown |
|  | 37 | 0 | *Picobirnaviridae* | unknown |
|  | 33 | 0 | *Hypoviridae* | unknown |
|  | 17 | 0 | unknown | unknown |
|  | 13 | 2 | *Adenoviridae* | *Mastadenovirus* |
|  | 13 | 2 | *Botourmiaviridae* | *Ourmiavirus* |
|  | 4 | 1 | *Chrysoviridae* | *Betachrysovirus* |
| C4 (FFPE: brain, liver, spleen) | 9063 | 0 | unknown | unknown |
|  | 2463 | 5 | *Chrysoviridae* | *Chrysovirus* |
|  | 1868 | 5 | *Botourmiaviridae* | *Ourmiavirus* |
|  | 1340 | 0 | *Partitiviridae* | unknown |
|  | 791 | 1 | *Partitiviridae* | *Gammapartitivirus* |
|  | 528 | 3 | *Chrysoviridae* | *Betachrysovirus* |
|  | 340 | 0 | *Picobirnaviridae* | unknown |
|  | 199 | 3 | *Arenaviridae* | *Reptarenavirus* |
|  | 18 | 1 | *Chrysoviridae* | *Alphachrysovirus* |
|  | 2 | 1 | *Endornaviridae* | *Alphaendornavirus* |
| C7 (cell culture medium) | 49006 | 132 | *Arenaviridae* | *Reptarenavirus* |
|  | 343 | 6 | *Arenaviridae* | *Hartmanivirus* |
| W1 (FFPE: liver) | 3570 | 7 | *Arenaviridae* | *Reptarenavirus* |
|  | 3 | 1 | *Adenoviridae* | *Mastadenovirus* |
| W2 (FFPE liver) | 10182 | 10 | *Arenaviridae* | *Reptarenavirus* |
| W3 (FFPE tissues) | 54831 | 8 | *Arenaviridae* | *Reptarenavirus* |
|  | 6 | 1 | Retroviridae | Gammaretrovirus |
| W4 (frozen liver) | 66140 | 200 | *Arenaviridae* | *Reptarenavirus* |
| W4 (cell culture medium) | 25399 | 18 | *Arenaviridae* | *Reptarenavirus* |
| W5 (frozen brain) | 29780 | 275 | *Arenaviridae* | *Reptarenavirus* |
|  | 1297 | 2 | *Arenaviridae* | *Hartmanivirus* |
| W5 (cell culture medium) | 23786 | 11 | *Arenaviridae* | *Reptarenavirus* |
|  | 298 | 7 | *Arenaviridae* | *Hartmanivirus* |

1. Plyusnin I, Kant R, Jääskeläinen AJ, Sironen T, Holm L, Vapalahti O, Smura T. 2020. Novel NGS pipeline for virus discovery from a wide spectrum of hosts and sample types. *Virus Evolution* 6.

The NGS approach led to identification of two pairs of hartmanivirus S and L segments, one in animal C7 and the other in animal W5. Pairwise sequence comparison (PASC) analysis (38) of the hartmanivirus segments found in animal C7 animal showed that the S segment has 75.61% nucleotide identity to Haartman Institute snake virus 1 (HISV-1) S segment and the L segment has 77.75% identity to the L segment of HISV-1. Similarly, the hartmanivirus S segment found in animal W5 67.06% identity to Dante Muikkunen virus 1 (DaMV-1) S segment whereas the L segment had 68.51% identity to the L segment of DaMV-1. We further compared the identified hartmanivirus segments to those available from GenBank by generating a similarity matrix, Table 3. According to the PASC analysis and the distance matrix, the hartmanivirus L segment identified from the samples of animal C7 could be classified into the same species with HISV-1 and −2 if strictly following the current arenavirus taxonomic demarcation criteria (9). However, the corresponding analyses for the S segment would support generation of a novel hartmanivirus species, due to which we named the corresponding virus as Universidad Nacional virus 1 (UnNV-1). Both PASC and similarity matrix analyses of the S and L segments identified in animal W5 supported the generation of a novel hartmanivirus species. We named the respective virus as big electron-dense squares virus (BESV-1) due to the presence of large square-shaped electron-dense accumulations observed in electron microscopy of BESV-1 infected cells. We further performed phylogenetic analysis of the hartmanivirus S and L segments (Fig. 6), which supports the classification of these viruses as novel hartmanivirus species. Specifically, on the basis of RdRp, UnNV-1 forms a sister clade for HISV-1 and HISV-2, whereas BESV-1 forms a sister clade for all DaMV-1, UnNV-1, HISV-1 and HISV-2 (Fig. 6A). Congruently with the similarity matrices, BESV-1 clustered together with DaMV-1 on the basis of GPC and NP and these two formed a sister clade for UnNV-1, HISV-1 and HISV-2 (Fig. 6B, C).

**Figure 6.**
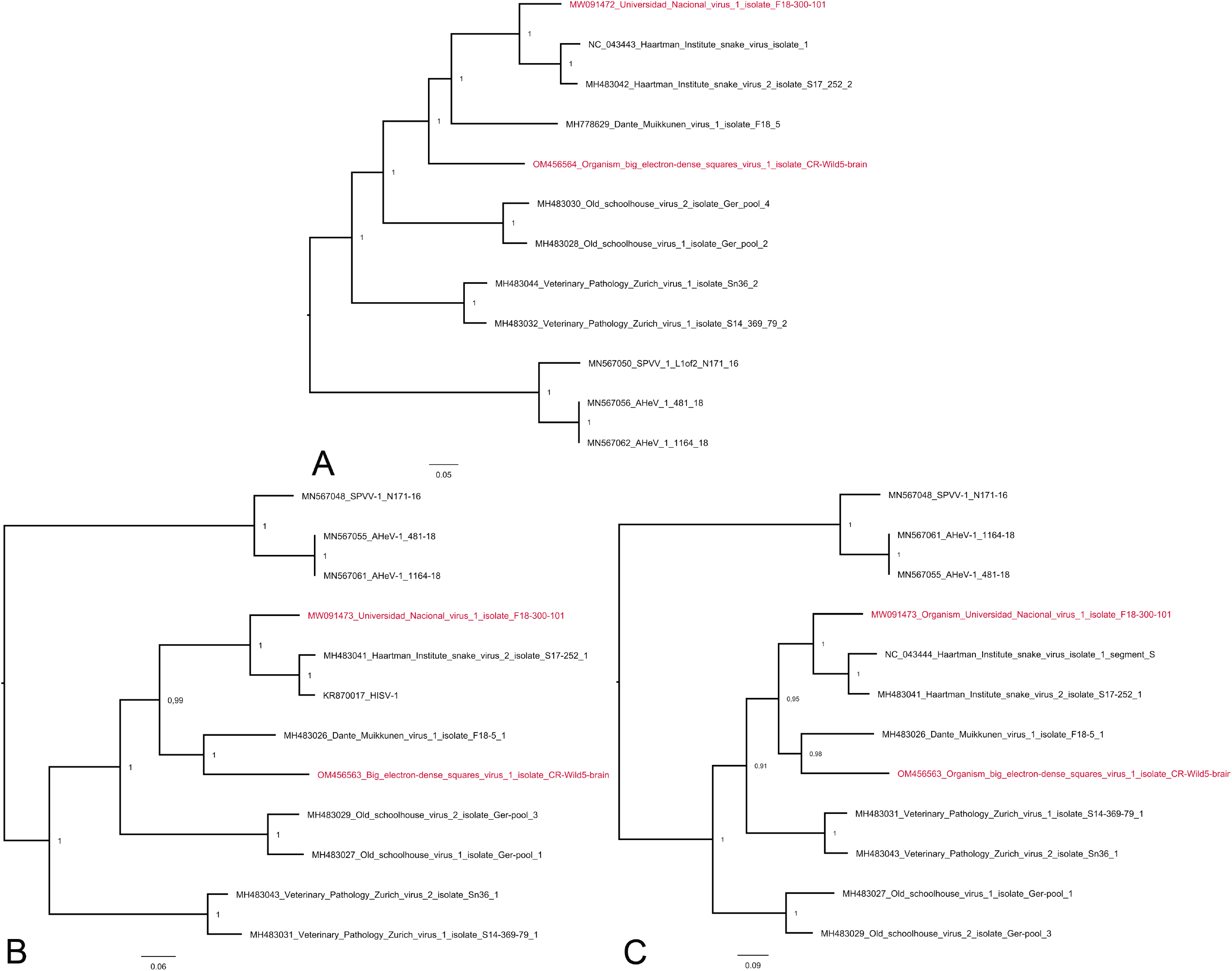
Phylogenetic analysis of hartmaniviruses identified in the study. The maximum clade credibility trees were inferred using the Bayesian MCMC method with Cprev Blosum and WAG amino acid substitution models for RdRp, GPC and NP, respectively. **A.** A phylogenetic tree based on the RdRp amino acid sequences of the viruses identified in this study and those available in GenBank. **B.** Phylogenetic tree based on the NP amino acid sequences of the viruses identified in this study and those available in GenBank. **C.** A phylogenetic tree based on the GPC amino acid sequences of the viruses identified in this study and those available in GenBank.

**Table 3.**
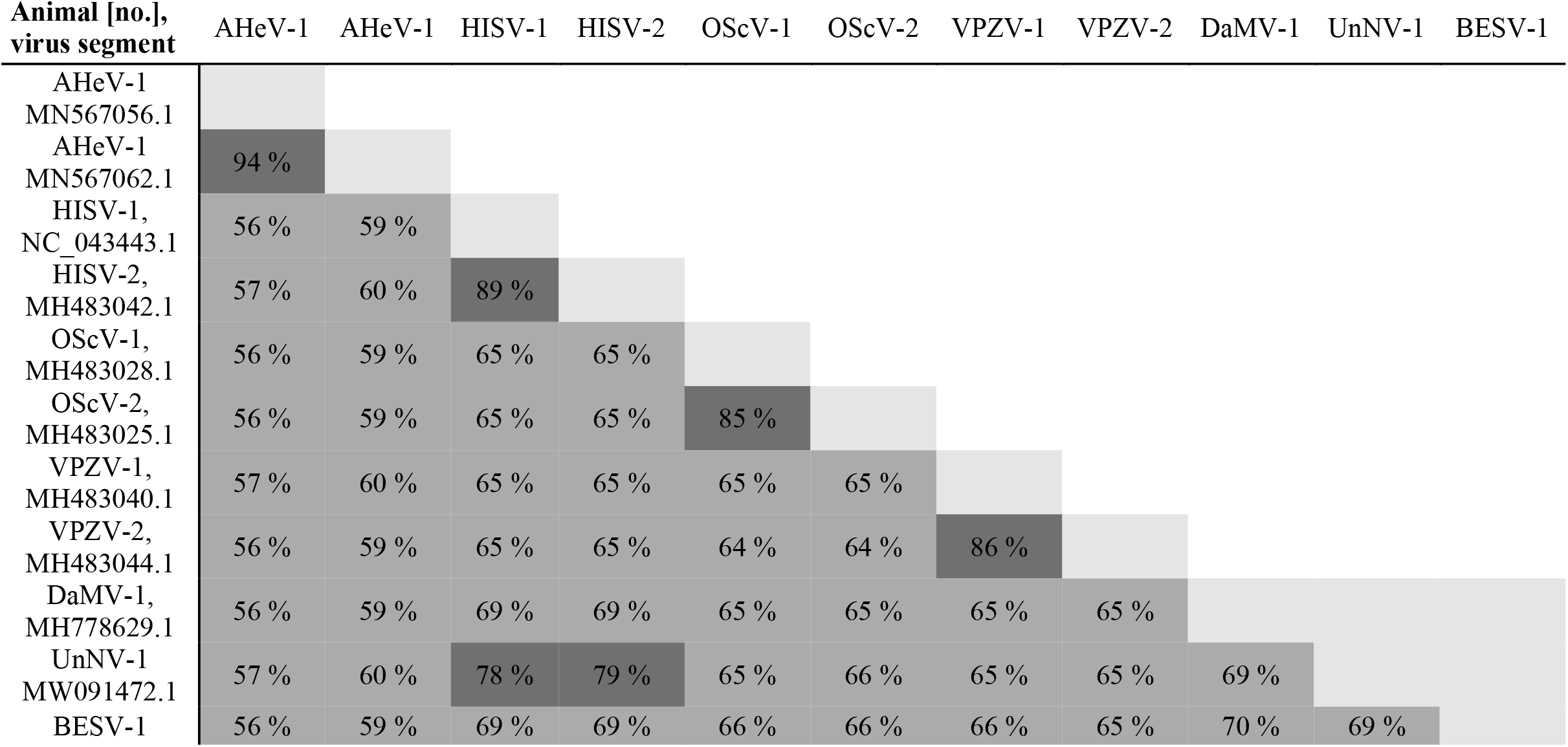
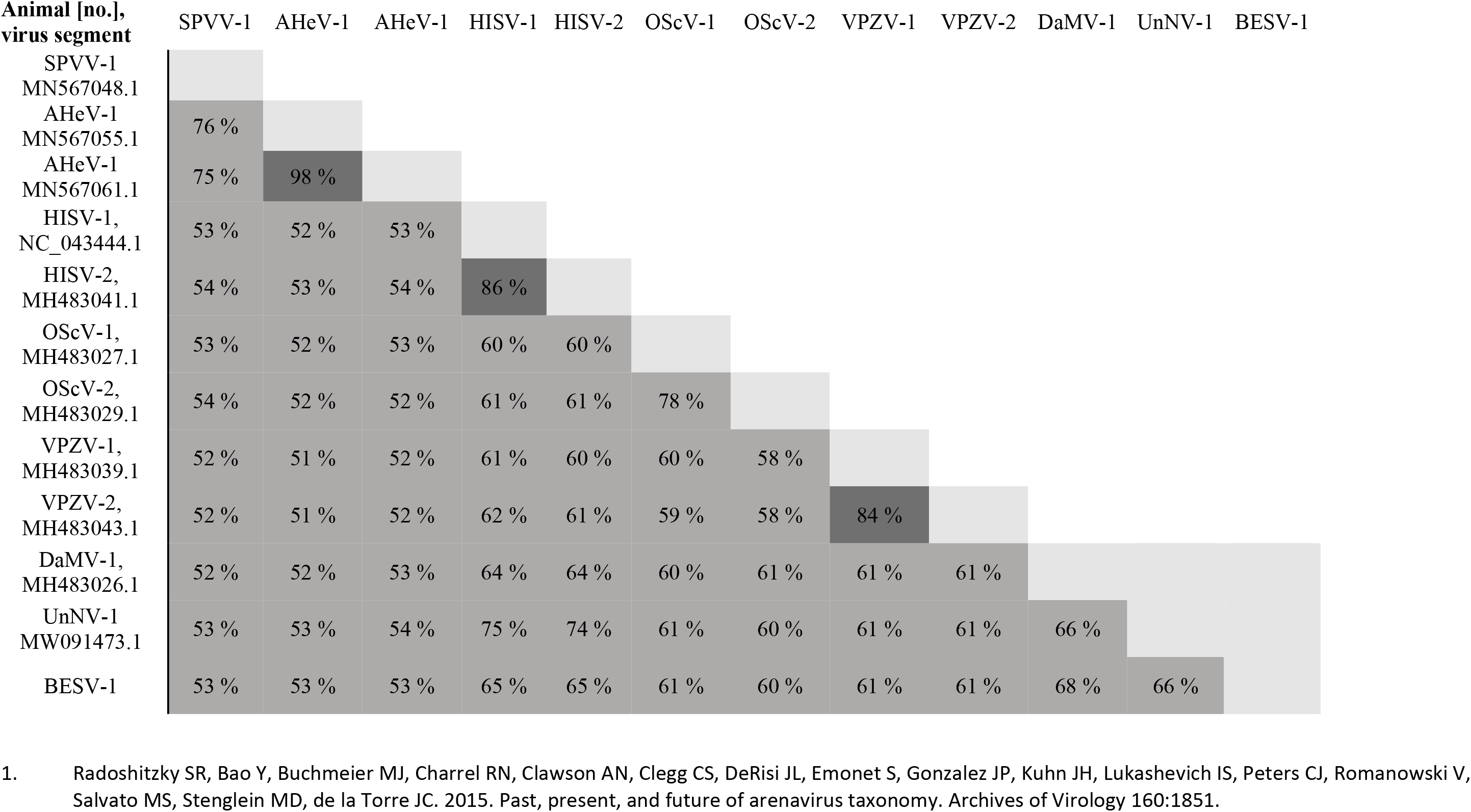
Comparison of hartmaniviruses identified this study to full-length sequences available from GenBank. **A)** Nucleotide identities between L segments. The numbers represent nucleotide identity, dark grey indicates >76% nucleotide identity (suggesting classification within same species, (1)). The abbreviations: AHeV=andere heimat virus, HISV=Haartman Institute snake virus, OScV=old schoolhouse virus, VPZV=veterinary pathology Zurich virus, DaMV=Dante Muikkunen virus, UnNV=Universidad Nacional virus, BESV=big electron-dense squares virus. **B)** Nucleotide identities between S segments. The numbers represent nucleotide identity, dark grey indicates >80% nucleotide identity (suggesting classification within same species, (1)). The abbreviations: AHeV=andere heimat virus, HISV=Haartman Institute snake virus, OScV=old schoolhouse virus, VPZV=veterinary pathology Zurich virus, DaMV=Dante Muikkunen virus, UnNV=Universidad Nacional virus, BESV=big electron-dense squares virus.

## DISCUSSION

The initial descriptions of BIBD date back to the 1970s and have only reported the disease in captive snakes (1). Prior to identification of reptarenaviruses as the causative agent (2-4, 7, 8), reports suggested that the disease can affect a multitude of snake species (1). The studies on reptarenaviruses have focused on captive constrictor snakes; it is so far unknown whether reptarenaviruses occur in wild animals and in different snake species as natural hosts. In a previous study, we provided evidence of reptarenavirus infections in native Brazilian boa constrictors by studying diseased snakes from wild animal sanctuaries, however, we could not rule out the possibility of reptarenavirus infection/transmission during captivity (10). Here, we report that BIBD occurs in indigenous captive snakes and wild boa constrictors in Costa Rica, and by studying archival tissue blocks demonstrate that the disease and its causative viruses have been present in wild *B. constrictor* snakes since at least the late 1980s.

The clinical signs and pathological findings in the wild boa constrictors are comparable to those of “classical” BIBD in the early descriptions of the disease in captive boas (11, 12). Interestingly, in the authors’ experience, similar findings have been rare among captive boas in Europe since the beginning of the new millennium. This could be a sign of adaptation of “the European reptarenavirome” to captive boa constrictor populations or rather that the variety of circulating L and S segments in European as compared to e.g. US captive collections is smaller. In the present study both wild and captive boa constrictors were found to have BIBD, and in *C. annulatus* the disease was restricted to captive individuals. This could suggest that the infection was transmitted to *C. annulatus*, directly or indirectly, from an infected boa or other animals of the collection.

Association of each mammarenavirus species with a specific rodent (or bat) host has led to the hypothesis of virus-host co-evolution (39). The current knowledge on reptarenaviruses presents a very different scenario in which the majority of snakes with BIBD carry several L and S segments of different reptarenavirus species (5, 6, 10, 16, 30). We cannot exclude that a sampling bias (examination of captive snakes with BIBD, and mainly boa constrictors) can explain the observed frequent co-infections. Snakes with BIBD often carry more L segments (we have identified up to seven in one snake, (30)) than S segments (we usually find 1 to 3 different S segments in one animal). Because the S segment encodes GPC, which gives rise to the spike complex that is essential for mediating the cell entry and fusion of the virion, one could speculate that the S segment may be under greater selection pressure. The spike complex that most efficiently mediates transmission and particle assembly would theoretically be enriched in the case of co-infection, assuming that L and S segments pair up rather freely as suggested by the data of Stenglein and colleagues (5). The L segment codes for the RdRp and the ZP proteins, and it seems reasonable to speculate that the RdRp would not distinguish between S (or L) segments of different virus species leading to transcription being equally effective for all segments.

With the above assumptions considered, one could speculate that the frequent reptarenaviruses co-infections seen in captive snakes (5, 6, 10, 16, 30) are a result of cross-species transmission of reptarenaviruses between snakes of different species and/or origin during transportation or co-housing. Reptarenavirus infection in boa constrictor is vertically transmitted both, from the father and the mother (30), which could have allowed the accumulation of L and S segment pools over a relatively long time (the first descriptions of BIBD are from the 1970s). The wild boa constrictors of this study appeared to carry only a single pair of L and S segments. Interestingly, the S segment found in these animals represents the same species as University of Giessen virus (UGV, or S6 segment following the nomenclature of Stenglein et al. (5)) which is at present the S segment most often found in captive boa constrictors with BIBD (5, 16, 30). If snakes were the reservoir hosts of reptarenaviruses, one could hypothesize that this particular S segment belongs to the reptarenaviruses that co-evolved with the boa constrictor. However, none of the animals included in our earlier study describing reptarenaviruses in native Brazilian boa constrictors carried the UGV-/S6-like S segment (10), thus negating the hypothesis. One could further hypothesize that the UGV-/S6-like segments seen in captive snakes in US and Europe would originate from wild-caught snakes in the Middle American region. The results also raise the general question as to whether a virus of which boas are the natural host would lead to BIBD, a disease that is eventually fatal, perhaps due to associated immunosuppression (1). If the reptarenaviruses found in the wild animals of this study are not native to boa constrictor, then how did the animals acquire the infection? Cross-species transmission from the prey animals of the snakes represent one option for acquiring the infection. The diet of wild boa constrictors in Costa Rica consists of a wide range of animals including birds, mammals, e.g. mice, rats, possums, bats, lizards, and iguanas. It is basically identical in Brazil, and different prey species could carry different reptarenaviruses, thus potentially explaining the wide variety of reptarenaviruses identified in snakes. However, using cultured cells of different animal species, we earlier demonstrated that reptarenavirus replication occurs at 30°C rather than at 37°C (40), which suggests the potential natural host to be cold-blooded/heterothermic. It would thus also be possible that boa constrictors acquire the infection through bites of blood-sucking parasites. In line with this arthropod transmission hypothesis, anecdotal evidence suggests that snake mite infestation of snake colonies coincides with the occurrence of BIBD (1). Furthermore, we have demonstrated that reptarenaviruses can infect cultured cells of three different tick species (40). Given the potential broad host range of the viruses, there is a need for further studies over a wide range of wild animals, including snakes, to identify the reservoir host of reptarenaviruses.

Alike reptarenaviruses, the origin of hartmaniviruses remains a mystery. We have identified several hartmaniviruses during our earlier NGS studies on snakes with BIBD (6, 16, 24, 35), however, unlike reptarenaviruses, hartmaniviruses appear to always be present as a single pair of S and L segments. So far, we have not associated hartmanivirus infection with any pathological findings or clinical signs, but neither have we been able to rule out their pathogenic potential. The present study identified two novel hartmaniviruses, UnNV-1 and BESV-1, and we managed to cultivate both viruses in boid kidney cells, albeit in co-infection with reptarenaviruses. Interestingly, the cultures with BESV-1 displayed large inclusions that were brightly eosinophilic in the hematoxylin-eosin stain. Ultrastructural analysis of the infected cells revealed the presence of large square-shaped electron-dense inclusions within the infected cells. The reptarenavirus S segment present in co-infection with BESV-1 hartmanivirus is classified into *Giessen reptarenavirus* species, the most commonly found reptarenavirus S segment in snakes with BIBD (5, 16). It is thus unlikely – and supported by our immunohistological findings - that the square-shaped inclusions observed in the infected cells do not associate with reptarenavirus NP expression. This could suggest that BESV-1 infection can induce IB formation, perhaps via NP expression, and would thus represent the first hartmanivirus associated with IB formation.

## ACKNOWLEDGEMENTS

The authors are grateful to laboratory staff of the Histology Laboratories, Departamento de Patologia, Escuela de Medicina Veterinaria, Universidad Nacional, Heredia Institute of Veterinary Pathology, and the Institute of Veterinary Pathology, Vetsuisse Faculty, University of Zürich, for excellent technical support. We also wish to acknowledge CSC – IT Center for Science, Finland, for computational resources. The study received financial support from the Leading House for the Latin American Region, University of St. Gallen, Switzerland, and the Academy of Finland (grant numbers 308613, 314119, and 335762).

## REFERENCES

1. Chang L-W, Jacobson ER. 2010. Inclusion Body Disease, A Worldwide Infectious Disease of Boid Snakes: A Review. Journal of Exotic Pet Medicine 19:216.

2. Stenglein MD, Sanders C, Kistler AL, Ruby JG, Franco JY, Reavill DR, Dunker F, Derisi JL. 2012. Identification, characterization, and in vitro culture of highly divergent arenaviruses from boa constrictors and annulated tree boas: candidate etiological agents for snake inclusion body disease. mBio 3:e00180.

3. Bodewes R, Kik MJ, Raj VS, Schapendonk CM, Haagmans BL, Smits SL, Osterhaus AD. 2013. Detection of novel divergent arenaviruses in boid snakes with inclusion body disease in The Netherlands. The Journal of general virology 94:1206.

4. Hetzel U, Sironen T, Laurinmaki P, Liljeroos L, Patjas A, Henttonen H, Vaheri A, Artelt A, Kipar A, Butcher SJ, Vapalahti O, Hepojoki J. 2013. Isolation, identification, and characterization of novel arenaviruses, the etiological agents of boid inclusion body disease. Journal of virology 87:10918.

5. Stenglein MD, Jacobson ER, Chang LW, Sanders C, Hawkins MG, Guzman DS, Drazenovich T, Dunker F, Kamaka EK, Fisher D, Reavill DR, Meola LF, Levens G, DeRisi JL. 2015. Widespread recombination, reassortment, and transmission of unbalanced compound viral genotypes in natural arenavirus infections. PLoS pathogens 11:e1004900.

6. Hepojoki J, Salmenpera P, Sironen T, Hetzel U, Korzyukov Y, Kipar A, Vapalahti O. 2015. Arenavirus Coinfections Are Common in Snakes with Boid Inclusion Body Disease. Journal of virology 89:8657.

7. Stenglein MD, Sanchez-Migallon Guzman D, Garcia VE, Layton ML, Hoon-Hanks LL, Boback SM, Keel MK, Drazenovich T, Hawkins MG, DeRisi JL. 2017. Differential Disease Susceptibilities in Experimentally Reptarenavirus-Infected Boa Constrictors and Ball Pythons. Journal of virology 91:10.1128/JVI.00451.

8. Hetzel U, Korzyukov Y, Keller S, Szirovicza L, Pesch T, Vapalahti O, Kipar A, Hepojoki J. 2021. Experimental Reptarenavirus Infection of Boa constrictor and Python regius. Journal of virology 95:e01968.

9. Radoshitzky SR, Bao Y, Buchmeier MJ, Charrel RN, Clawson AN, Clegg CS, DeRisi JL, Emonet S, Gonzalez JP, Kuhn JH, Lukashevich IS, Peters CJ, Romanowski V, Salvato MS, Stenglein MD, de la Torre JC. 2015. Past, present, and future of arenavirus taxonomy. Archives of Virology 160:1851.

10. Argenta FF, Hepojoki J, Smura T, Szirovicza L, Hammerschmitt ME, Driemeier D, Kipar A, Hetzel U. 2020. Identification of Reptarenaviruses, Hartmaniviruses and a Novel Chuvirus in Captive Brazilian Native Boa Constrictors with Boid Inclusion Body Disease. Journal of virology doi:JVI.00001-20 [pii].

11. Schumacher J, Jacobson ER, Homer BL, Gaskin JM. 1994. Inclusion Body Disease in Boid Snakes. Journal of Zoo and Wildlife Medicine 25:511.

12. Wozniak E, McBride J, DeNardo D, Tarara R, Wong V, Osburn B. 2000. Isolation and Characterization of an Antigenically Distinct 68- kd Protein from Nonviral Intracytoplasmic Inclusions in Boa Constrictors Chronically Infected with the Inclusion Body Disease Virus (IBDV: Retroviridae). Veterinary Pathology Online 37:449.

13. Keilwerth M, Buhler I, Hoffmann R, Soliman H, El-Matbouli M. 2012. Inclusion Body Disease (IBD of Boids)--a haematological, histological and electron microscopical study. Berliner und Munchener tierarztliche Wochenschrift 125:411.

14. Chang LW, Fu A, Wozniak E, Chow M, Duke DG, Green L, Kelley K, Hernandez JA, Jacobson ER. 2013. Immunohistochemical detection of a unique protein within cells of snakes having inclusion body disease, a world-wide disease seen in members of the families Boidae and Pythonidae. PloS one 8:e82916.

15. Chang L, Fu D, Stenglein MD, Hernandez JA, DeRisi JL, Jacobson ER. 2016. Detection and prevalence of boid inclusion body disease in collections of boas and pythons using immunological assays. Veterinary journal (London, England : 1997) 218:13.

16. Windbichler K, Michalopoulou E, Palamides P, Pesch T, Jelinek C, Vapalahti O, Kipar A, Hetzel U, Hepojoki J. 2019. Antibody response in snakes with boid inclusion body disease. PloS one 14:e0221863.

17. Hyndman TH, Marschang RE, Bruce M, Clark P, Vitali SD. 2019. Reptarenaviruses in apparently healthy snakes in an Australian zoological collection. Australian Veterinary Journal 97:93.

18. Turchetti A, Tinoco H, de Campos Cordeiro Malta M, Loyola Teixeira da Costa E, Pessanha A, Soave S, Paixao A, Santos R. 2013. Inclusion body disease in a Corallus hortulanus. 6:15.

19. Solórzano AL. 2004. Serpientes de Costa Rica: distribución, taxonomía e historia natural. INBio.

20. Henderson RW, Pauers MJ, Colston TJ. 2013. On the congruence of morphology, trophic ecology, and phylogeny in Neotropical treeboas (Squamata: Boidae: Corallus). Biological Journal of the Linnean Society 109:466–475.

21. de Pinho GM, de Lima DO, da Costa PN, F.A. dS. 2009. Natural history notes: Epicrates cenchria (Brazilian Rainbow Boa). Diet. Herpetological Review 40(3):354–355.

22. Radoshitzky SR, Buchmeier MJ, Charrel RN, Clegg JCS, Gonzalez JJ, Gunther S, Hepojoki J, Kuhn JH, Lukashevich IS, Romanowski V, Salvato MS, Sironi M, Stenglein MD, de la Torre JC, Ictv Report C. 2019. ICTV Virus Taxonomy Profile: Arenaviridae. The Journal of general virology 100:1200.

23. Abudurexiti A, Adkins S, Alioto D, Alkhovsky SV, Avsic-Zupanc T, Ballinger MJ, Bente DA, Beer M, Bergeron E, Blair CD, Briese T, Buchmeier MJ, Burt FJ, Calisher CH, Chang C, Charrel RN, Choi IR, Clegg JCS, de la Torre JC, de Lamballerie X, Deng F, Di Serio F, Digiaro M, Drebot MA, Duan X, Ebihara H, Elbeaino T, Ergunay K, Fulhorst CF, Garrison AR, Gao GF, Gonzalez JJ, Groschup MH, Gunther S, Haenni AL, Hall RA, Hepojoki J, Hewson R, Hu Z, Hughes HR, Jonson MG, Junglen S, Klempa B, Klingstrom J, Kou C, Laenen L, Lambert AJ, Langevin SA, Liu D, Lukashevich IS, et al. 2019. Taxonomy of the order Bunyavirales: update 2019. Archives of Virology 164:1949.

24. Hepojoki J, Hepojoki S, Smura T, Szirovicza L, Dervas E, Prahauser B, Nufer L, Schraner EM, Vapalahti O, Kipar A, Hetzel U. 2018. Characterization of Haartman Institute snake virus-1 (HISV-1) and HISV-like viruses-The representatives of genus Hartmanivirus, family Arenaviridae. PLoS pathogens 14:e1007415.

25. Maes P, Alkhovsky SV, Bao Y, Beer M, Birkhead M, Briese T, Buchmeier MJ, Calisher CH, Charrel RN, Choi IR, Clegg CS, de la Torre JC, Delwart E, DeRisi JL, Di Bello PL, Di Serio F, Digiaro M, Dolja VV, Drosten C, Druciarek TZ, Du J, Ebihara H, Elbeaino T, Gergerich RC, Gillis AN, Gonzalez JJ, Haenni AL, Hepojoki J, Hetzel U, Ho T, Hong N, Jain RK, Jansen van Vuren P, Jin Q, Jonson MG, Junglen S, Keller KE, Kemp A, Kipar A, Kondov NO, Koonin EV, Kormelink R, Korzyukov Y, Krupovic M, Lambert AJ, Laney AG, LeBreton M, Lukashevich IS, Marklewitz M, Markotter W, et al. 2018. Taxonomy of the family Arenaviridae and the order Bunyavirales: update 2018. Archives of Virology doi:10.1007/s00705-018-3843-5 [doi].

26. Aqrawi T, Stohr AC, Knauf-Witzens T, Krengel A, Heckers KO, Marschang RE. 2015. Identification of snake arenaviruses in live boas and pythons in a zoo in Germany. Tierarztliche PraxisAusgabe K, Kleintiere/Heimtiere 43:239.

27. Simard J, Marschang RE, Leineweber C, Hellebuyck T. 2020. Prevalence of inclusion body disease and associated comorbidity in captive collections of boid and pythonid snakes in Belgium. PloS one 15:e0229667.

28. Reed RNR, G.H. 2009. Giant constrictors: Biological and management profiles and an establishment risk assessment for nine large species of pythons, anacondas, and the boa constrictor, vol Open-File Report 2009–1202. U.S. Geological Survey

29. Reynolds RG, Henderson RW. 2018. Boas of the World (Superfamily Booidae): A Checklist With Systematic, Taxonomic, and Conservation Assessments. Bulletin of the Museum of Comparative Zoology 162:1–58, 58.

30. Keller S, Hetzel U, Sironen T, Korzyukov Y, Vapalahti O, Kipar A, Hepojoki J. 2017. Co-infecting Reptarenaviruses Can Be Vertically Transmitted in Boa Constrictor. PLoS pathogens 13:e1006179.

31. Baggio F, Hetzel U, Nufer L, Kipar A, Hepojoki J. 2020. Arenavirus nucleoprotein localizes to mitochondria. bioRxiv doi:10.1101/2020.11.06.370825:2020.11.06.370825.

32. Dervas E, Hepojoki J, Laimbacher A, Romero-Palomo F, Jelinek C, Keller S, Smura T, Hepojoki S, Kipar A, Hetzel U. 2017. Nidovirus-Associated Proliferative Pneumonia in the Green Tree Python (Morelia viridis). Journal of virology doi:JVI.00718-17 [pii].

33. Katoh K, Standley DM. 2013. MAFFT Multiple Sequence Alignment Software Version 7: Improvements in Performance and Usability. Molecular biology and evolution 30:772.

34. Ronquist F, Teslenko M, van der Mark P, Ayres DL, Darling A, Hohna S, Larget B, Liu L, Suchard MA, Huelsenbeck JP. 2012. MrBayes 3.2: efficient Bayesian phylogenetic inference and model choice across a large model space. Systematic Biology 61:539.

35. Hetzel U, Szirovicza L, Smura T, Prahauser B, Vapalahti O, Kipar A, Hepojoki J. 2019. Identification of a Novel Deltavirus in Boa Constrictors. mBio 10:10.1128/mBio.00014.

36. Plyusnin I, Kant R, Jääskeläinen AJ, Sironen T, Holm L, Vapalahti O, Smura T. 2020. Novel NGS pipeline for virus discovery from a wide spectrum of hosts and sample types. Virus Evolution 6.

37. Langmead B, Salzberg SL. 2012. Fast gapped-read alignment with Bowtie 2. Nature methods 9:357.

38. Bao Y, Chetvernin V, Tatusova T. 2014. Improvements to pairwise sequence comparison (PASC): a genome-based web tool for virus classification. Archives of Virology 159:3293.

39. Zapata JC, Salvato MS. 2013. Arenavirus variations due to host-specific adaptation. Viruses 5:241.

40. Hepojoki J, Kipar A, Korzyukov Y, Bell-Sakyi L, Vapalahti O, Hetzel U. 2015. Replication of Boid Inclusion Body Disease-Associated Arenaviruses Is Temperature Sensitive in both Boid and Mammalian Cells. Journal of virology 89:1119.

